# The Chaperone NASP Contributes to *De Novo* Deposition of the Centromeric Histone Variant CENH3 in *Arabidopsis* Early Embryogenesis

**DOI:** 10.1101/2023.10.05.560999

**Authors:** Hidenori Takeuchi, Shiori Nagahara, Tetsuya Higashiyama, Frédéric Berger

**Author notes:** Correspondence: Hidenori Takeuchi.

## Abstract

The centromere is an essential chromosome region where the kinetochore is formed to control equal chromosome distribution during cell division. The centromere-specific histone H3 variant CENH3 (also called CENP-A) is a prerequisite for the kinetochore formation. Since CENH3 evolves rapidly, associated factors, including histone chaperones mediating the deposition of CENH3 on the centromere, are thought to act through species-specific amino-acid sequences. The functions and interaction networks of CENH3 and histone chaperons have been well-characterized in animals and yeasts. However, molecular mechanisms involved in recognition and deposition of CENH3 are still unclear in plants. Here, we used a swapping strategy between domains of CENH3 of *Arabidopsis thaliana* and the liverwort *Marchantia polymorpha* to identify specific regions of CENH3 involved in targeting the centromeres and interacting with the general histone H3 chaperone, NASP (nuclear autoantigenic sperm protein). CENH3’s LoopN-α1 region was necessary and sufficient for the centromere targeting in cooperation with the α2 region and was involved in interaction with NASP in cooperation with αN, suggesting a species-specific CENH3 recognition. In addition, by generating an *Arabidopsis nasp* knockout mutant in the background of a fully fertile *GFP-CENH3*/*cenh3-1* line, we found that NASP was implicated for *de novo* CENH3 deposition after fertilization and thus for early embryo development. Our results imply that the NASP mediates the supply of CENH3 in the context of the rapidly evolving centromere identity in land plants.

## Introduction

In nuclei of eukaryotic cells, DNA is organized into chromatin that consists of an array of nucleosomes and associated proteins. The nucleosome, a basic unit of chromatin, is formed with ∼150 bp DNA and four core histones, H2A, H2B, H3, and H4 (Luger et al. 1997). Most eukaryote species possess a set of variants for each core histone, so-called histone variants, and use them to confer specific properties on each genomic region (Pusarla and Bhargava 2005, Talbert and Henikoff 2010, Kawashima et al. 2015, Jiang and Berger 2017). The histone monomer consists of the histone-fold domain, which contains α-helices connected by loops and mediates histone-histone and histone-DNA interaction, and the unstructured N- and C-terminal tails, which protrude from the nucleosome core particle and generally undergo covalent modifications, including methylation, acetylation, phosphorylation, and ubiquitination (Luger et al. 1997, Khorasanizadeh 2004).

The histone variants H3.1 and H3.3 are incorporated into nucleosomes of non-centromeric chromatin during DNA replication and throughout cell cycle, respectively (Filipescu et al. 2013, Talbert and Henikoff 2017, Borg et al. 2021). The histone variant CENH3 specifically marks the centromere. Due to the importance of CENH3 targeting to the centromere in eukaryotes, regulatory factors and DNA sequences related to CENH3 deposition as well as their evolution have been extensively studied in yeast, animals, and plants (Talbert et al. 2002, Malik et al. 2002, Henikoff and Dalal 2005, Morris and Moazed 2007, Zhang et al. 2008, Malik and Henikoff 2009, Hirsch et al. 2009, Nagaki et al. 2010, Yuan et al. 2015, Maheshwari et al. 2017, Rosin and Mellone 2017, Naish et al. 2021). In eukaryotes, the deposition of CENH3 proteins, also called CENP-A in mammals, CID in flies, Cse4 in budding yeast *Saccharomyces cerevisiae*, and Cnp1 in fission yeast *Schizosaccharomyces pombe*, is required for proper point kinetochore formation and thus equal chromosome distribution to daughter cells after cell division. In contrast with other H3 variants, the amino-acid sequences of CENH3 proteins are variable among species, even close relatives (Malik et al. 2002, Nagaki et al. 2010, Sanei et al. 2011, Rosin and Mellone 2016). Similarly, both centromeric DNA repeats and centromeric/heterochromatic proteins evolve rapidly.

In contrast with relatively-conserved canonical H3s, rapidly evolving CENH3 proteins are recognized by distantly-related or non-conserved histone chaperones, HJURP in mammals, Scm3 in yeasts, and CAL1 in flies (Camahort et al. 2007, Stoler et al. 2007, Dunleavy et al. 2009, Foltz et al. 2009, Zhou et al. 2011, Phansalkar et al. 2012, Chen et al. 2014). In humans and yeasts, the chaperones HJURP and Scm3 recognize the loop 1 and α2 helix (designated as the CENP-A targeting domain, CATD) within the histone-fold domain of CENH3 (CENP-A/Cse4) (Cho and Harrison 2011, Bassett et al. 2012). In *Drosophila* species, CAL1 distinguishes *Drosophila* CENH3 homologs via the loop 1 in a species-specific manner (Rosin and Mellone 2016). HJURP, Scm3, and CAL1 chaperones specifically interact with CENH3 and mediate *de novo* CENH3 deposition via self-sustaining epigenetic mechanism, which maintains centromere positioning and occasionally creates novel centromeres (Barnhart et al. 2011, Roure et al. 2019, Medina-Pritchard et al. 2020, Palladino et al. 2020). Similar to these features of animals and yeasts, it is suggested in plants that CENH3 recognition depends on its species-specific sequence and that CENH3 deposition is mediated by a yet uncharacterized self-sustaining mechanism (Ravi et al. 2010, Maheshwari et al. 2015, Marimuthu et al. 2021). In flowering plants (angiosperms), failure of CENH3 deposition and function leads to uniparental genome elimination after fertilization and consequently to generation of haploid individuals, which is utilized as doubled haploid technology to rapidly obtain homozygous lines (Ravi and Chan 2010, Ravi et al. 2014). Therefore, molecular mechanisms involved in CENH3 recognition and deposition have been intensively studied in crop species as well as *A. thaliana* (Lermontova et al. 2006, Ingouff et al. 2010, Sanei et al. 2011, Karimi-Ashtiyania et al. 2015, Kuppu et al. 2020). However, our knowledge about the requirements for recognition and deposition of plant CENH3 is still limited.

A homolog of NASP (nuclear autoantigenic sperm protein) in *A. thaliana* is also characterized as a common histone H3 chaperone (Maksimov et al. 2016, Le Goff et al. 2020). NASP is a tetratricopeptide repeat (TPR)-containing protein conserved in wide-range of eukaryotes and is involved in various cellular events, such as DNA replication, cell proliferation, cell growth, and embryonic development in mammals (Richardson et al. 2000, 2006, Dunleavy et al. 2007, Finn et al. 2012). NASP binds to H3-H4 dimers to reserve them in the soluble/nonnucleosomal fraction and hand off them to chromatin assembly factors, suggesting the function of histone supply chain for the demand in mammalian cells (Campos et al. 2010, Cook et al. 2011). Human NASP protein possess the direct nucleosome assembly activity toward CENH3 (CENP-A) as well as the canonical H3.1 and H3.3 *in vitro* (Osakabe et al. 2010). In *S. pombe*, NASP (called as Sim3) binds to CENH3 (Cnp1) and is required for its deposition on centromere (Dunleavy et al. 2007). Recent studies in *A. thaliana* have identified NASP as an interactor of both H3.3 and CENH3 through co-immunoprecipitation and mass spectrometry analysis using *A. thaliana* tissues or cultured cell line (Maksimov et al. 2016, Le Goff et al. 2020). Similar to yeast and human NASPs, *A. thaliana* NASP is broadly expressed in actively dividing tissues and is a soluble protein localized at the nucleoplasm. *In vitro* experiments using purified histones and NASP have demonstrated that *A. thaliana* NASP binds monomeric H3.1 and H3.3 as well as H3.1-H4 and H3.3-H4 dimers, but not monomeric H4, stabilizes H3-H4 tetramers from H3-H4 dimers, and promotes tetrasome formation with DNA (Maksimov et al. 2016). Interaction assay using a bimolecular fluorescence complementation (BiFC) have suggested that *A. thaliana* NASP interacts with both N-terminal tail and histone-fold domain of CENH3 *in planta* (Le Goff et al. 2020). Knockdown of *NASP* expression causes moderate reduction of nuclear CENH3 levels and ∼30% reduction of CENH3 signal at the centromere (Le Goff et al. 2020).

Here we show large-scale and detailed structure-function analysis of plant CENH3 in terms of centromere targeting in *A. thaliana* cells and investigate NASP function in *A. thaliana* embryogenesis by generating a *nasp* knockout mutant. By a domain swapping strategy using CENH3 proteins of *A. thaliana* and the liverwort *Marchantia polymorpha*, we reveal the requirement of specific CENH3 regions, such as LoopN-α1 region, for centromere targeting and interaction with NASP in a lineage-specific manner. Additionally, we characterize the role of NASP in *de novo* CENH3 deposition after fertilization

## Results and Discussion

### Targeting *A. thaliana* centromere by CENH3 protein involves its angiosperm-specific amino-acid sequence context

To understand a lineage-specific recognition mechanism of plant CENH3 proteins for the centromere targeting, we compared CENH3 amino-acid sequences of the dicot *A. thaliana* (HTR12; AtCENH3), the monocot *Zea mays* (ZmCENH3) that has previously shown to target *A. thaliana* centromere (Ravi et al. 2010), the liverwort *M. polymorpha* (MpCENH3), and the human (CENP-A) (Fig. 1A and Supplementary Fig. S1). In contrast to the canonical H3.3 histones, which share the same amino-acid sequences between *A. thaliana* and *M. polymorpha* and have only six amino-acid differences between human and land plants (Supplementary Fig. S1A), CENH3 proteins are highly variable (Fig. 1A) as illustrated previously (Ravi et al. 2010). The N-terminal tail of CENH3 proteins is especially hypervariable with different length, in contrast with the strong degree of conservation of the histone-fold domain (Fig. 1A and Supplementary Fig. S1B).

**Fig. 1.**
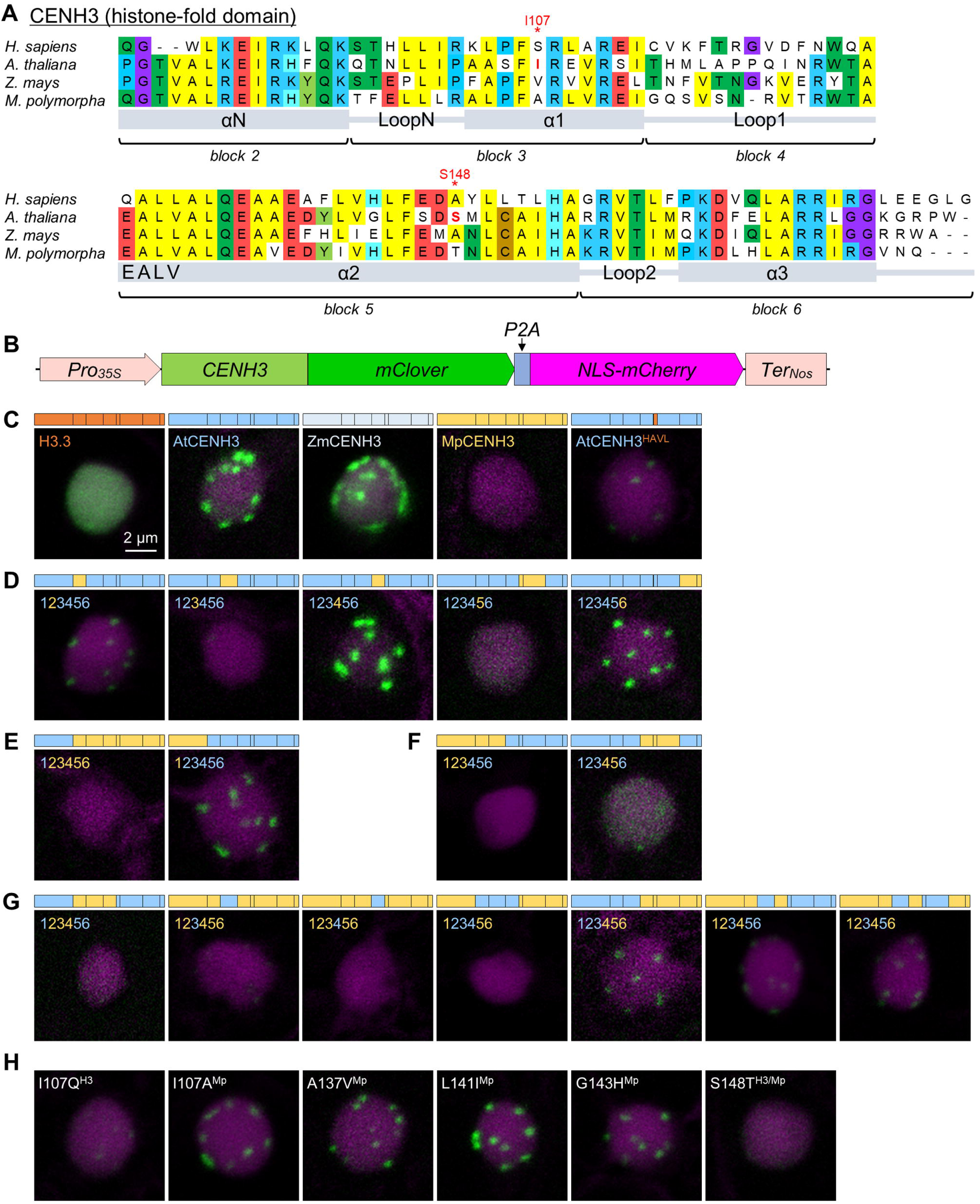
CENH3 protein regions required for targeting *A. thaliana* centromere. (A) Sequence alignment of histone-fold domain of *H. sapiens*, *A. thaliana*, *Z. mays*, and *M. polymorpha* CENH3 proteins. Protein secondary structure and blocks dividedly analyzed in this study are shown below the alignment. See also Supplementary Figure S1A and B for sequence alignments of the canonical H3 and N-terminal tail of CENH3. (B) Schematic of the expression construct to produce CENH3 variants fused to mClover and NLS-mCherry from a single transcript. Co-translational “self-cleavage” during translation of the 2A peptide sequence (*P2A*) leads to generate two separate proteins. (C-H) Localization assessment of CENH3 variants in the *A. thaliana* petal cell. Block bars drawn above each image and titles guide the protein structure of each variant with colors. Each figure panel consisting of multiple variants represents comparisons which examine canonical H3 (H3.3), CENH3 of *A. thaliana* (AtCENH3), *Z. mays* (ZmCENH3), and *M. polymorpha* (MpCENH3), and AtCENH3 substituted with HAVL (AtCENH3^HAVL^) (C); necessary single AtCENH3 region (D); importance of N-terminal tail (E) and CATD (F); sufficient AtCENH3 region (G); and necessary amino acid of AtCENH3 (H). The images are representative nuclei from more than three images capturing multiple nuclei. See also Supplementary Figure 2 and 3 for summary of this assay and representative images of similar assay using tobacco BY-2 cells.

Previous studies have shown that CENH3 proteins from angiosperms, including several Brassicaceae species, the grapevine *Vitis vinifera*, and the monocot *Z. mays*, are able to target *A. thaliana* centromeres and to complement the embryo lethality of *cenh3-1* null mutation when expressed without GFP tag (Ravi et al. 2010, Maheshwari et al. 2015). The histone-fold domain of CENH3 is sufficient to target the centromere (Ravi and Chan 2010). To obtain further insight into the sequence specificity for the CENH3 function, we used the MpCENH3 sequence of the distant land plant *M. polymorpha*, which is substantially different from *A. thaliana* CENH3 in the histone-fold domain [58% (56/97), including gaps], even though ZmCENH3 is similarly different (58%, 57/98) (Fig. 1A). To assess the protein localization of CENH3 variants after virtually comparable translation levels in plant cells, we constructed an expression vector that consists of the *cauliflower mosaic virus 35S* promoter, *mClover* encoding the green fluorescent protein, and *NLS-mCherry* for the nuclear-localized red fluorescent protein, connected by *P2A* sequence for the 2A peptide from porcine teschovirus-1 (Fig. 1B). The P2A sequence causes co-translational “self-cleavage” during translation as a result of ribosome skipping (Kawashima et al. 2014; Liu et al. 2017), which generates mClover-fused CENH3 variant and NLS-mCherry proteins from a single mRNA.

To obtain steady and reliable results, we generated transgenic *A. thaliana* individuals harboring series of CENH3 variants and observed mClover and mCherry fluorescence in the petal, which contains actively dividing cells and is useful to capture fluorescent signals with lower background (Fig. 1C-H). In NLS-mCherry-positive nuclei, AtCENH3-mClover and ZmCENH3-mClover marked dot-like centromere regions. As a control, the exchange of the four contiguous amino acids EALV located N-terminal of α2 helix following Loop1, which distinguish CENH3, to HAVL from H3.3 caused a dramatic reduction of the mClover signal with only faint dot-like localization (Fig. 1C, AtCENH3^HAVL^). Contrasting with AtCENH3 and ZmCENH3, MpCENH3-mClover signal were undetectable (Fig. 1C), suggesting that MpCENH3 misses an angiosperm-specific sequence necessary for its deposition on the centromeres.

### CENH3 deposition on *A. thaliana* centromeres requires its LoopN-**α**1 region

We next sought to define specific regions of CENH3 important for the lineage-specific amino-acid recognition in the *A. thaliana* cell. By generating a series of transgenic *A. thaliana* expressing chimeric CENH3 proteins, which consist of each block from AtCENH3 and MpCENH3 (Fig. 1A and Supplementary Fig. S2). Swapping one block of AtCENH3 by that of MpCENH3 indicated that LoopN-α1 (block 3) and α2 (block 5) were required for targeting *A. thaliana* centromere (Fig. 1D). The N-terminal tail from AtCENH3 was not required for centromere targeting of MpCENH3 (Fig. 1E), although the N-terminal tail is important for full CENH3 functions in *A. thaliana* (Ravi and Chan 2010, Maheshwari et al. 2015). In human cells, a domain spanning Loop1 to α2 (designated as CATD) of CENP-A conferred centromere targeting and function when swapped into canonical human H3 (Black et al. 2007). In contrast, a corresponding domain of AtCENH3 failed to complement *A. thaliana cenh3-1* phenotype when swapped into H3.3 (Ravi et al. 2010). We also assessed the impact of CATD (block 4 and 5) of AtCENH3 on localization in *A. thaliana* cell by using MpCENH3 as a donor. The CATD and a larger domain containing the CATD and the α3 region (block 6) of AtCENH3 were not sufficient for centromere targeting (Fig. 1F and Supplementary Fig. S2). On the contrary, mClover signal of a chimeric AtCENH3 having CATD of MpCENH3 was detected in the nucleus and weakly focused on dot-like regions resembling centromeres (Fig. 1F and Supplementary Fig. S2), suggesting that regions outside of CATD are involved in escort and reserve of plant CENH3 at the nucleoplasm through lineage-specific amino-acid recognition.

To determine sufficient regions for *A. thaliana* centromere targeting in the plant CENH3 framework, we analyzed further chimeric CENH3 variants (Fig. 1G). In the presence of the LoopN-α1 (block 3) of AtCENH3, additional AtCENH3 regions, such as N-terminal tail and αN regions (block 1-2) or α2 region (block 5), conferred the capability to target centromeres in the lineage-specific manner. We searched angiosperms’ CENH3-specific amino acids responsible for targeting *A. thaliana* centromere and found that two AtCENH3 mutants possessing single amino acid substitution, I107Q and S148T, abolished centromere targeting (Fig. 1H).

To complement these results obtained in the *A. thaliana* cell, we also observed tobacco BY-2 cultured cell lines harboring each chimeric CENH3 construct (Supplementary Fig. S2 and S3). We obtained fundamentally similar results in the BY-2 system to the *A. thaliana* system: LoopN-α1 (block 3) of AtCENH3 was the necessary region for centromere targeting. The I107A mutant, which has a single substitution by MpCENH3-type amino acid, abolished specific targeting to the centromere in the BY-2 nucleus, while the S148T mutant was localized at centromere at comparable level to wild-type AtCENH3 (Supplementary Fig. S3). In summary, by the swapping strategy using AtCENH3 and MpCENH3 sequences, we found that the CENH3’s LoopN-α1 region in combination with additional regions, such as α2, is the key for the centromere targeting in *A. thaliana* and BY-2 cells and indicated the lineage-specific amino-acid recognition mechanism of plant CENH3.

### The general H3/CENH3 chaperone NASP contributes to embryogenesis in *A. thaliana*

In most eukaryotes, CENH3 is the indispensable protein by the common function in centromere specification to form the kinetochore, whereas it evolves rapidly. In accordance with this divergence, distantly-related or non-conserved histone chaperones, HJURP in mammals, Scm3 in yeasts, and CAL1 in flies have been identified as species-specific chaperons for CENH3/CENP-A/Cse4/CID (Stoler et al. 2007, Camahort et al. 2007, Dunleavy et al. 2009, Foltz et al. 2009, Zhou et al. 2011, Phansalkar et al. 2012, Chen et al. 2014). Although no specific chaperone that specifically deposit CENH3 have been identified in the plant lineage, one of the homologs of histone chaperones found in animals and yeasts, NASP, has been reported to be a general H3 chaperone that escorts CENH3 as well as H3.1 and H3.3 into the nucleoplasm in *A. thaliana* (Maksimov et al. 2016, Le Goff et al. 2020).

We initially tried to identify a novel specific chaperone that functions in a species-specific CENH3 recognition by our reverse-genetic mini-screening of CENH3-related factors, including unknown function genes co-expressed with *CENH3*, but we failed to get putative knockout mutants showing the embryo lethality similar to *cenh3* mutant. We also generated CRISPR/Cas9-mediated mutants for homologs of histone chaperones in the transgenic plant line *GCH3*, that expresses *GFP-CENH3* in *cenh3-1* mutant background (Ravi et al. 2010). We obtained a heterozygous 10-bp deletion mutant of *NASP* gene in line *GCH3* (*nasp^cri^/NASP GCH3*) but never obtained a homozygous *nasp^cri^ GCH3* mutant from progeny of the heterozygous line. In self-pollinated *nasp^cri^/NASP GCH3* siliques approximately 25% of developing seeds were white embryoless seeds or collapsed brown seeds, indicative of the embryonic defect (Fig. 2A-D). About 22% of ovules in self-pollinated *nasp^cri^/NASP GCH3* siliques had an arrested embryo at 1- to 2-cell stages, compared with ovules of wild-type Col-0 and *GCH3* containing a normal heart stage embryo (Fig. 2E, F). This phenotype was not observed after genetic complementation in T1 plants carrying the *NASPpro::NASP-mRuby2* (native promoter-driven genomic *NASP* fused with red fluorescent protein gene *mRuby2*) (Fig. 2D, *At^genomic^*). From these results, we concluded that the 10-bp deletion recessive mutation was responsible for the loss of function NASP and the observed embryogenesis defect in homozygous *nasp^cri^ GCH3* seeds. Our results first indicate that *NASP* is indispensable in the *A. thaliana* development in a certain condition, at least *GCH3* background.

**Fig. 2.**
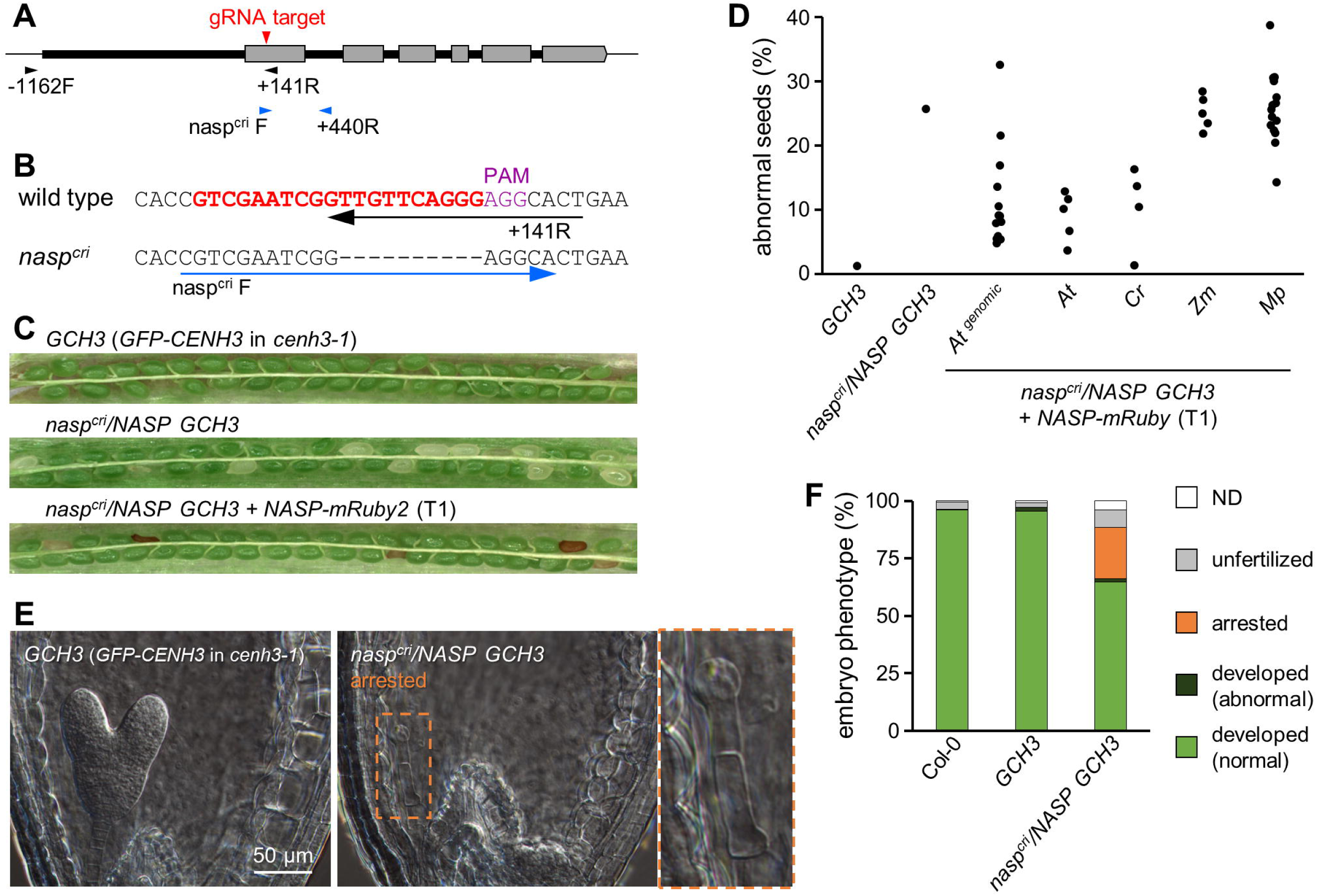
Embryo lethal phenotype of *nasp^cri^* knockout mutant in *GCH3* line. (A) Schematic of *NASP* gene structure with a guide RNA (gRNA) target site and genotyping primers. Gray boxes are exons of *NASP* gene. The bold line indicates a region used for complementation constructs. (B) Primer design to exclusively amplify endogenous *NASP* gene and *nasp^cri^* mutant allele. (C) Representative images of developing seeds in pistils of a *GCH3* background line (*GFP-CENH3* in *cenh3-1*), a heterozygous *nasp^cri^/NASP* in *GCH3*, and a complemented line with *NASP-mRuby2* transgene. (D) The percentage of abnormal seeds per developing seeds in self-pollinated siliques. For T1 lines, siblings from a *nasp^cri^/NASP GCH3* plant were used for the introduction of each orthologous *NASP* gene fused to *mRuby2*, and T1 plants with *nasp^cri^/NASP* genotype were selected by genomic PCR. Each dot represents the sum value of five siliques for each independent T1 plant. (E) Embryo development in ovules from *GCH3* or *nasp^cri^/NASP GCH3* five days after pollination. A heart-stage embryo in *GCH* ovule and an early arrested embryo at 1-cell stage in self-pollinated *nasp^cri^/NASP GCH3* ovule. (F) The proportion of embryo phenotype. The phenotype was categorized based on observation as follows: developed (normal), about heart-stage embryo; developed (abnormal), globular to heart-stage embryo with distorted shape; arrested, 1- or 2-cell stage embryo; unfertilized, non-elongated egg cell or no egg cell structure due to degeneration; ND, not defined. 209–321 ovules/seeds from 4 to 6 siliques were analyzed for each genotype.

### NASP is required for early embryo development after the first zygotic division of the *A. thaliana GFP-CENH3* line

To clarify the contribution of NASP to embryogenesis and CENH3 deposition after fertilization, we obtained *GCH3* plant lines with homozygous *nasp^cri^* mutation complemented with hemizygous *NASPpro::NASP-mRuby2* transgene. By observing red fluorescence of NASP-mRuby2 at the nucleus, we were able to distinguish whether zygotes and early embryos in self-pollinated pistils consisted of cells of functional *NASP* (homozygous or hemizygous *NASPpro::NASP-mRuby2*) or *nasp^cri^ GCH3* mutant (Fig. 3). At 2 days after pollination (2 DAP), most NASP-mRuby2-positive embryos normally developed at 4- or 8-cell stages, whereas NASP-mRuby2-negative ones were at 1- or 2-cell stages (Fig. 3A, C). At 4 DAP, NASP-mRuby2-positive embryos further developed at about the early heart stage without any obvious morphological defects, whereas NASP-mRuby2-negative ones were still at the 1- or 2-cell stages (Fig. 3B). Normally developing NASP-mRuby2-positive embryo cells exhibited GFP-CENH3 signals, while the NASP-mRuby2-negative embryo cells had no GFP-CENH3 signals in the nucleus (Fig. 3A, B). These results suggested that the absence of NASP caused failure of CENH3 deposition after fertilization causing the early arrest of the embryo at the 1- or 2-cell stages.

**Fig. 3.**
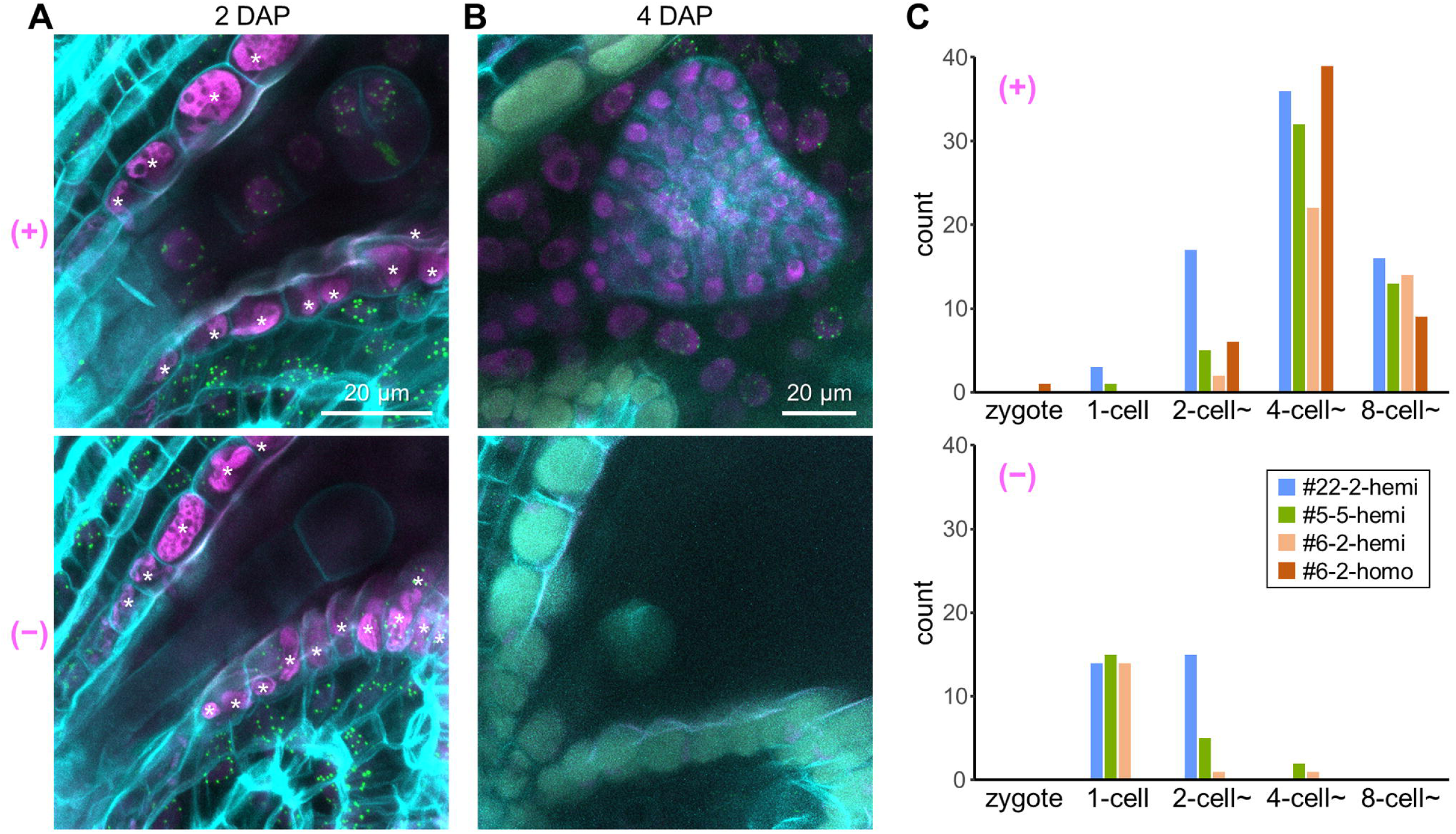
Requirement of NASP for early embryogenesis and *de novo* CENH3 deposition in *GCH* line. (A, B) Confocal images of ClearSee-treated ovules inside the pistil of homozygous *nasp^cri^ GCH3* with hemizygous *NASPpro::NASP-mRuby2* transgene at 2 days after pollination (A) and 4 days after pollination (B). The presence (+) or absence (−) of mRuby2 red fluorescence (shown as magenta) in embryo nuclei indicate NASP-complemented embryo or homozygous *nasp^cri^* mutant embryo, respectively. The GFP-CENH3 signals at centromeres were observed in the embryo and endosperm of NASP-mRuby2-positive (+) ovules and in the maternal integuments of ovules regardless of NASP-mRuby2 existence in embryo cells (+ and −). Note that asterisks indicate unrelated objectives with autofluorescence in ovular integument cells. Green, GFP-CENH3; cyan, cell walls stained with calcofluor white. (C) The number of ovules with each stage of zygote/embryo from three independent hemizygous lines and one homozygous line for *NASPpro::NASP-mRuby2* transgene in homozygous *nasp^cri^ GCH3*. Counting was conducted using confocal images of ClearSee-treated ovules (*n* = 54–101) from 3 siliques at 2 days after pollination.

### Species-specific sequences of NASP involve its function in embryogenesis

Taking advantage of the *nasp^cri^/NASP GCH3* line that showed the clear embryogenesis phenotype (Fig. 2C, E), we assessed the importance of NASP protein and its domains in terms of species-specific amino-acid sequences by the complementation test. As similar to the assessment for CENH3 as shown in Fig. 1, we focused on NASP from *A. thaliana*, a Brassicaceae species *Capsella rubella*, *Z. mays*, and *M. polymorpha*. The introduction of coding sequences for AtNASP and CrNASP but not ZmNASP and MpNASP restored the abnormal seed phenotype of *nasp^cri^/NASP GCH3* (Fig. 2D), suggesting that NASP required a certain degree of species-specific amino-acid sequence for its function.

We compared amino-acid sequences of NASP from human and these four plant species (Fig. 4A). NASP proteins include four tetratricopeptide repeat (TPR) motifs and a C-terminal extension containing a nuclear localization signal (Dunleavy et al. 2007, Bowman et al. 2015, Liu et al. 2021, Bao et al., 2022, Liu et al. 2022). The NASPs contains an acidic amino acid-rich stretch as an interrupted loop in the second TPR, which seems to be variable between species, and a less conserved N-terminal region without any predicted protein structures (Fig. 4A). To examine the requirement of each NASP region for the function in *A. thaliana* embryogenesis, we introduced AtNASP variants fused with the cyan fluorescent protein mTurquoise2 into *nasp^cri^/NASP GCH3*. Compared with full-length wild-type AtNASP (*At*) as a control, AtNASP lacking the N-terminal region [*At* (Δ*N*)] complemented the abnormal seed phenotype despite lower frequency and degree, and AtNASP lacking the acidic region [*At* (Δ*DE*)] did not at all (Fig. 4B). Some T1 line of AtNASP having N-terminal region of MpNASP [*At* (*N^Mp^*)] just slightly decreased the abnormal seed phenotype, while more than half of T1 lines of AtNASP having the acidic region of MpNASP [*At* (*DE^Mp^*)] restored the phenotype as wild-type AtNASP (Fig. 4B). Consistent of this, the abnormal embryo phenotype was restored by the introduction of *At* (*DE^Mp^*) but not *At* (*N^Mp^*) or MpNASP (*Mp*) (Fig. 4C). This analysis suggested that the acidic region was required for NASP function. Our observations are also consistent with the importance of the acidic region of human NASP for the binding to H3 to make the complex with H3–H4–ASF1 (Bowman et al. 2017).

**Fig. 4.**
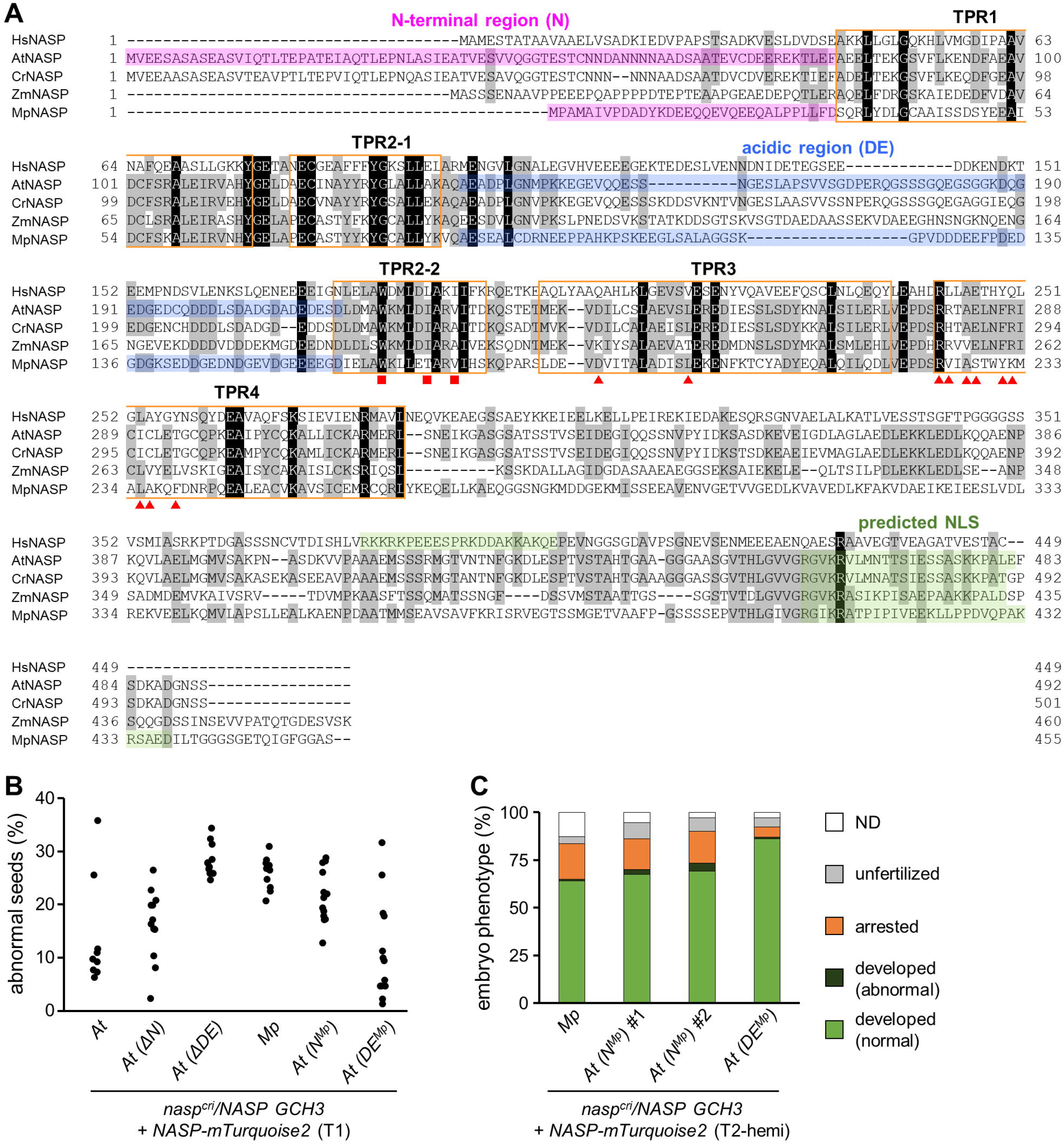
NASP protein domains required for the function in *A. thaliana* embryo development. (A) Sequence alignment of NASP proteins of *H. sapiens* (HsNASP), *A. thaliana* (AtNASP), *Z. mays* (ZmNASP), and *M. polymorpha* (MpNASP). Red squares and triangles below the alignment indicate amino acids of AtNASP that has been shown to mediate binding to canonical H3’s NLterminal and α3 regions, respectively (Liu et al. 2022). See also Supplementary Figure S1C and D for sequence alignments of these binding regions of H3 in comparison with AtCENH3 and MpCENH3. (B) The percentage of abnormal seeds per developing seeds in self-pollinated siliques from T1 lines. As same as Fig. 2D, siblings from a *nasp^cri^/NASP GCH3* plant were used for the introduction of each *NASP* variant fused to *mTurquoise2*, and T1 plants with *nasp^cri^/NASP* genotype were selected by genomic PCR. Each dot represents the sum value of five siliques for each independent T1 plant. (C) The proportion of embryo phenotype. The phenotype was categorized as noted for Fig. 2F. 101–246 ovules/seeds from 3 to 7 siliques were analyzed for each genotype.

### *A. thaliana* NASP interacts with CENH3 through lineage-specific sequence contexts

Our complementation test implied the requirement of lineage-specific sequence contexts for the NASP function. Since amino-acid sequences of canonical histone H3 are completely the same among land plant species (Supplementary Fig. S1A), we postulated that the lineage-specific NASP-CENH3 interaction would be one of explanations for the functionality of NASP variants. To assess whether CENH3 variants, including chimera of AtCENH3 and MpCENH3, interacted with AtNASP, we performed semi-quantitative interaction assay by a bimolecular fluorescence complementation (BiFC) assay in *N. benthamiana* leaf epidermal cells. As a control, we also assessed the interaction between CENH3 variants and *A. thaliana* ASF1A, a conserved histone chaperone for H3/H4 (Zhu et al. 2011, Min et al. 2019, Zhong et al. 2022).

In our BiFC assay employing the P2A-NLS-mCherry (PNC) module as used in Fig. 1, YFP fluorescence (BiFC signal) should be observed if AtASF1A or AtNASP fused to nYFP interacted with H3.3 or CENH3 variants fused to cYFP-PNC, while mCherry fluorescence certifies comparable translation level of H3.3 or CENH3 variants (Fig. 5). The BiFC assay indicated that AtASF1A interacted with both AtCENH3 and MpCENH3 but AtNASP did only with conspecific AtCENH3 (Fig. 5A). AtASF1A interacted with H3.3, ZmCENH3, and all of At/Mp-chimeric CENH3 variants we tested, despite slightly different relative BiFC intensity (Fig. 5B). In contrast, AtNASP interacted with AtCENH3, ZmCENH3, and some chimeric CENH3 variants but not the others as well as MpCENH3 (Fig. 5C). When a single region, LoopN-α1 (block 3) or α2 (block 5) of AtCENH3, which were required for the targeting to *A. thaliana* centromere (Fig. 1D), was swapped by that of MpCENH3, these chimeric CENH3 variants retained the interaction capability with AtNASP (Fig. 5C_i). The introduction of neither block 3 nor blocks 3/5 of AtCENH3 conferred the interaction capability to MpCENH3 framework (Fig. 5C_ii). Interestingly, CENH3 variants with blocks 1/2/3 or blocks 2/3/4 of AtCENH3 interacted with AtNASP, while that of MpCENH3 did not (Fig. 5C_iii). This clear contrast demonstrated that the NASP-CENH3 interaction relied on angiosperm-specific sequence contexts of CENH3’s αN and LoopN-α1. Together with the result that the LoopN-α1 region of AtCENH3 played the dominant role in the centromere targeting of *A. thaliana* and tobacco BY-2 cells (Fig. 1, Supplementary Fig. 2 and 3), our data suggest that NASP could recognize CENH3 through the LoopN-α1 region and escort it into the nucleoplasm for the deposition on the centromere nucleosome.

**Fig. 5.**
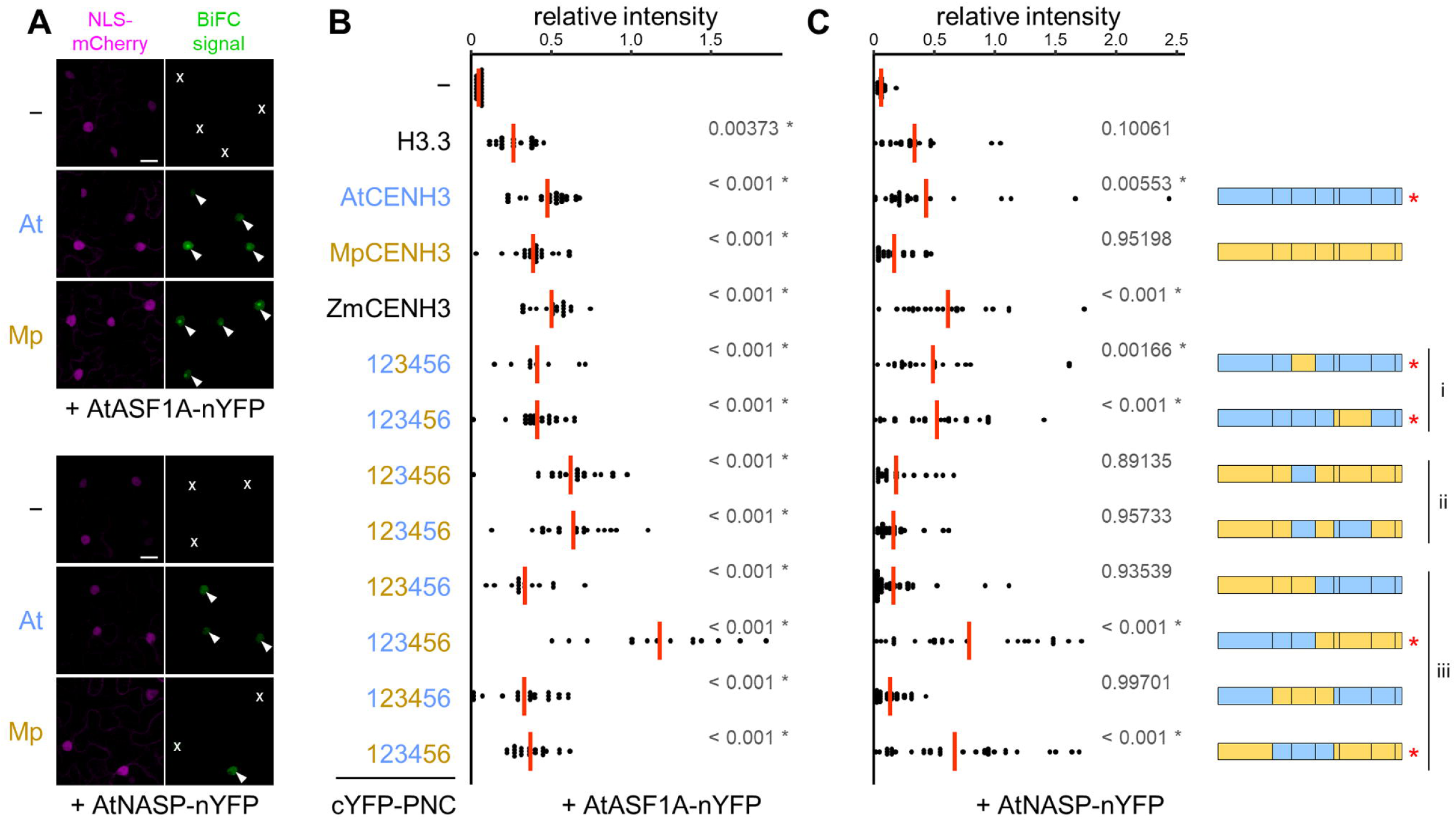
Interaction assay for CENH3 variants and AtASF1A or AtNASP using a BiFC system with P2A-NLS-mCherry (PNC) module. (A) Confocal images of *N. benthamiana* leaf epidermal cells of BiFC assay between AtASF1A-nYFP or AtNASP-nYFP and cYFP-PNC (−), AtCENH3-cYFP-PNC (At), or MpCENH3-cYFP-PNC (Mp). Equivalent NLS-mCherry signal implies comparable levels of the three cYFP fusion proteins. Nuclei with negative or positive BiFC signals were marked with x signs or arrowheads, respectively. Scale bars, 20 µm. (B, C) The relative BiFC intensity (BiFC/mCherry) for H3.3, AtCENH3, MpCENH3, ZmCENH3, or At/Mp-chimeric CENH3 variants with AtASF1A (B) and AtNASP (C). Each dot represents the value of each nucleus. Red lines show mean of the values. *P* values shown in gray are estimated by the Dunnett’s test compared with negative control (−). For AtCENH3, MpCENH3, and At/Mp-chimeric CENH3 variants, block bars on the right indicate the structure of each variant with colors (light blue, AtCENH3; yellow, MpCENH3), and red asterisks indicate CENH3 variants showing significant BiFC signal.

A previous study using structural and biochemical approaches for AtNASP and canonical H3 showed that AtNASP has distinct two sets of amino-acids clusters for the binding to H3’s N□terminal (a.a. 21–59 containing αN) and α3 (a.a. 116–135) regions, respectively, *in vitro* (Liu et al. 2022, Fig. 4A, Supplementary Fig. S1C, D), similar to the binding mode of human NASP to H3 (Bowman et al. 2017, Liu et al. 2021, Bao et al., 2022). Although this knowledge is the case for canonical H3, AtNASP might also bind to CENH3 via these regions, which are not completely conserved between AtCENH3 and MpCENH3 (Supplementary Fig. S1C, D). Since our interaction assay using *N. benthamiana* implied that AtNASP required AtCENH3’s LoopN-α1 in addition to αN, but not α3, for the species-specific NASP-CENH3 interaction (Fig. 5C), the recognition of CENH3 by NASP could be mediated by different dynamics *in planta*. Altogether, our results indicated that NASP acts as a key chaperone that bindCENH3 in a species-specific manner in plants. It should be noted that the completely-arrested embryonic phenotype by the loss-of-function *nasp* mutation was only observed in the *GCH3* background, so the GFP tag of CENH3 might interfere interaction with some other components important for normal CENH3 supply network, including another general H3 chaperone ASF1 (Fig. 5B, Cook et al. 2011).

## Conclusions

In this study, we defined *A. thaliana* CENH3’s domains important for centromere targeting and identified NASP as a chaperone that binds to CENH3 through a domain around LoopN-α1 in a species-specific sequence manner. In addition, we demonstrated that NASP is involved in early embryo development and CENH3 deposition after fertilization. Although NASP seems to be neither a CENH3-specific chaperone nor a sole chaperone that recognizes CENH3, we propose the hypothesis that NASP contributes to *de novo* deposition of CENH3 on the centromere after fertilization. This function requires species-specific sequences, suggesting coevolution of NASP with CENH3 after fertilization. Our findings help to understand how the CENH3 protein is deposited at the centromere in the zygote nucleus and contribute to haploid induction technology by manipulation of CENH3 deposition network.

## Materials and Methods

### Plant materials

The *A. thaliana* Col-0 was used for the expression and localization experiment of CENH3 variants. The *GFP-CENH3* in *cenh3-1* (*GCH3*) line was kindly provided by Dr. Simon Chan’s laboratory (Ravi et al. 2010). For transformation of *A. thaliana* plants, T-DNA constructs were introduced by floral dip using *Agrobacterium tumefaciens* strain GV3101. *A. thaliana* plants were germinated on half-strength Murashige-Skoog (MS) solid medium containing antibiotics (if applicable) under continuous light at 22°C and transferred on soil for further growth under long-day condition in plant growth rooms. The *N. benthamiana* plant was grown from directly sowing seeds on soil in plant growth rooms under long-day condition at 22-26°C. Transformation of the tobacco BY-2 cultured cell was performed as previously described (Mayo et al. 2006) using *A. tumefaciens* strain EHA105. Several transformants per each construct were selected via several round of subculturing of callus on modified LS medium containing 1.5% agar, 50 mg/L kanamycin and 100 mg/L cefotaxime sodium. Each BY-2 callus was observed by suspending in liquid modified LS medium.

### Plasmids and sequences

All plasmids were generated by standard molecular techniques and listed in Supplementary Table S1. The *P2A-NLS-mCherry* sequence was amplified from a plasmid obtained from Dr. Tomokazu Kawashima (Kawashima et al. 2014). The *nasp^cri^* mutant was generated by CRISPR/Cas9 system based on pKIR1.1 (Tsutsui and Higashiyama 2017). Gene and protein sequence data can be obtained from The Arabidopsis Information Resource (TAIR; http://arabidopsis.org) for *A. thaliana* and from Phytozome (https://phytozome-next.jgi.doe.gov/) for *C. rubella*, *Z. mays*, and *M. polymorpha*.

### Confocal microscopy and image analysis

Confocal images of petal cells, ovules, and BY-2 cells of transgenic lines were taken using an LSM-780-DUO-NLO system (Zeiss). Images were acquired using ZEN 2010 software (Zeiss) and processed using using Fiji software (http://fiji.sc/ Fiji).

### Establishment of the *nasp^cri^* mutant line

T1 seeds transformed with HTv985, pKIR1.1 vector containing guide RNA (gRNA) sequence (5′-GTCGAATCGGTTGTTCAGGG-3′) that targets the first exon of *NASP* gene (Fig. 2A), were selected by red fluorescence of seeds. One mutant candidate (#3) was obtained by genomic PCR using primers [NASP+3F (5′-GGTTGAAGAATCAGCTTC-3′) and NASP+440R (5′-CATGATTAGATCATTCGAG-3′)] and direct Sanger sequencing, which showed a disrupted sequence around the gRNA target site. From seeds harvested from this candidate T1 individual, Cas9-free plants were selected based on the absence of red fluorescence of seeds and confirmed later by no hygromycin resistant seedling in the next generation. Genomic PCR and sequencing analysis identified the 10-bp deletion in the *nasp^cri^/NASP* heterozygous plant (#3-16). The siblings from #3-16 plant were used for phenotypic analysis and transformation for complementation tests.

Based on the 10-bp deletion, NASP+141R (5′-AGTGCCTCCCTGAACAAC-3′) for wild-type *NASP* and nasp^cri^ F (5′-CGTCGAATCGGAGGCA-3′) for 10-bp deletion *nasp^cri^* allele were designed (Fig. 2B). Primers for wild type allele [NASP-1162F (5′-AATACTAAGCGAGCCATC-3′) and NASP+141R] and primers for *nasp^cri^* allele (nasp^cri^ F and NASP+440R) were used to exclusively amplify endogenous *NASP* gene and *nasp^cri^* allele, respectively.

### Evaluation of embryo phenotype by clearing in chloral hydrate solution

The carpel walls were removed from siliques five days after hand pollination using tweezers and 27-gauge needle (Terumo, Tokyo, Japan). The remaining tissues, ovules/seeds attached to the septum, were fixed and decolorized in ethanol/acetic acid (9:1) and rehydrated with ethanol series (80%, 60%, 40%, and 20%) and distilled water.

After rehydration, the samples were cleared in chloral hydrate solution (chloral hydrate, glycerol, and distilled water in the weight ratio of 8:1:3). Differential interference contrast (DIC) microscopy analysis of cleared ovules/seeds was performed with a Nikon Ts2R microscope.

### Evaluation of embryo development with fluorescence by ClearSee

Two or four days after hand pollination, siliques were dissected as described above and fixed by immersing in 4% paraformaldehyde/PBS and applying a reduced pressure with a vacuum pump. About 1 hour after the fixation, samples were washed with PBS twice and cleared with ClearSeeAlpha solution to reduce the formation of brown pigment in the ClearSee-treated ovule (Kurihara et al. 2021). 2–3 weeks after treatment in the dark at room temperature, the samples were stained with ClearSee solution (Kurihara et al. 2015) containing 0.1 mg/mL calcofluor white for 1 hour and then washed with ClearSee solution for 1 hour. The silique sample was split at the center of the septum to separate two lines of placenta with seeds into one line of that, which facilitates the observation of embryo phenotype without overlapping each seed tissue. For confocal observation using LSM-780-DUO-NLO system (Zeiss), images were acquired with a 40× objective lens (LD C-Apochromat 40x/1.1 W Corr, Zeiss) under 488/561 nm excitation and 490-561/588-695 nm emission wavelengths for mClover and mRuby2 florescence and sequentially under 405 nm excitation and 415-490 nm emission wavelength for calcofluor white.

### BiFC assay

Transient expression in *N. benthamiana* leaves by agroinfiltration was performed as previously described (Takeuchi and Higashiyama 2016). Equal amounts of *A. tumefaciens* strain GV3101 cultures for nYFP and cYFP-PNC constructs and the p19 silencing suppressor were collected in a tube and resuspended in infiltration buffer (10 mM MgCl_2_, 10 mM MES, pH 5.6, and 150 μM acetosyringone). Three to four hours after incubation at 26°C, the mixed suspensions were infiltrated in four to five-week-old *N. benthamiana* leaves. Two days after infiltration, the leaf samples were subjected to confocal microscope observation.

To virtually normalize protein production levels among CENH3 variants fused to cYFP-PNC sequence, the relative BiFC intensity was BiFC signal (YFP fluorescence) divided by NLS-mCherry signal. For each channel of images, background was subtracted and fluorescent intensity of nuclei region were measured. For each BiFC combination, the relative intensity of 9–36 nuclei from 3 replicate samples was calculated and plotted using R studio (https://cran.ism.ac.jp/). Statistical analysis was also performed using R studio (“multcomp” package; https://CRAN.R-project.org/package=multcomp) based on the Dunnett’s multiple comparison test.

## Data Availability

The data underlying this study, including nucleotide and protein sequences, are available in the article and supplementary data.

## Acknowledgments

We thank Simon Chan’s laboratory for *GFP-CENH3 in cenh3-1* seed; Hiroki Tsutsui for pKIR1.1 vector; Tomokazu Kawashima for plasmids and cDNAs used for plasmid construction; Akihisa Osakabe and other members of Berger lab (GMI, Vienna, Austria) for helpful discussion; Daisuke Kurihara for technical assistance of BY-2 experiment; Minako Ueda for technical advice on observation of zygote and early embryo; and Yoshikatsu Sato and Nagoya University Live Imaging Center for Zeiss LSM-780-DUO-NLO microscope system.

## Author Contributions

H.T. conceived and designed this study under supervision of F.B.; H.T. generated the research materials; H.T. and S.N. performed the experiments and analyzed the data; H.T. wrote the manuscript with input from S.N., T.H., and F.B.

## References

Bao, H., Carraro, M., Flury, V., Liu, Y., Luo, M., Chen, L., et al. (2022) NASP maintains histone H3-H4 homeostasis through two distinct H3 binding modes. Nucleic Acids Res. 50: 5349–5368.

Barnhart, M.C., Kuich, P.H.J.L., Stellfox, M.E., Ward, J.A., Bassett, E.A., Black, B.E., et al. (2011) HJURP is a CENP-A chromatin assembly factor sufficient to form a functional de novo kinetochore. J Cell Biol. 194: 229–243.

Bassett, E.A., DeNizio, J., Barnhart-Dailey, M.C., Panchenko, T., Sekulic, N., Rogers, D.J., et al. (2012) HJURP Uses Distinct CENP-A Surfaces to Recognize and to Stabilize CENP-A/Histone H4 for Centromere Assembly. Dev Cell. 22: 749–762.

Black, B.E., Jansen, L.E.T., Maddox, P.S., Foltz, D.R., Desai, A.B., Shah, J. V., et al. (2007) Centromere Identity Maintained by Nucleosomes Assembled with Histone H3 Containing the CENP-A Targeting Domain. Mol Cell. 25: 309–322.

Borg, M., Jiang, D., and Berger, F. (2021) Histone variants take center stage in shaping the epigenome. Curr Opin Plant Biol. 61.

Bowman, A., Koide, A., Goodman, J.S., Colling, M.E., Zinne, D., Koide, S., et al. (2017) sNASP and ASF1A function through both competitive and compatible modes of histone binding. Nucleic Acids Res. 45: 643–656.

Bowman, A., Lercher, L., Singh, H.R., Zinne, D., Timinszky, G., Carlomagno, T., et al. (2015) The histone chaperone sNASP binds a conserved peptide motif within the globular core of histone H3 through its TPR repeats. Nucleic Acids Res. 44: 3105–3117.

Camahort, R., Li, B., Florens, L., Swanson, S.K., Washburn, M.P., and Gerton, J.L. (2007) Scm3 Is Essential to Recruit the Histone H3 Variant Cse4 to Centromeres and to Maintain a Functional Kinetochore. Mol Cell. 26: 853–865.

Campos, E.I., Fillingham, J., Li, G., Zheng, H., Voigt, P., Kuo, W.H.W., et al. (2010) The program for processing newly synthesized histones H3.1 and H4. Nat Struct Mol Biol. 17: 1343–1351.

Chen, C.C., Dechassa, M.L., Bettini, E., Ledoux, M.B., Belisario, C., Heun, P., et al. (2014) CAL1 is the Drosophila CENP-A assembly factor. J Cell Biol. 204: 313–329.

Cho, U.S., and Harrison, S.C. (2011) Recognition of the centromere-specific histone Cse4 by the chaperone Scm3. Proc Natl Acad Sci U S A. 108: 9367–9371.

Cook, A.J.L., Gurard-Levin, Z.A., Vassias, I., and Almouzni, G. (2011) A Specific Function for the Histone Chaperone NASP to Fine-Tune a Reservoir of Soluble H3-H4 in the Histone Supply Chain. Mol Cell. 44: 918–927.

Dunleavy, E.M., Pidoux, A.L., Monet, M., Bonilla, C., Richardson, W., Hamilton, G.L., et al. (2007) A NASP (N1/N2)-Related Protein, Sim3, Binds CENP-A and Is Required for Its Deposition at Fission Yeast Centromeres. Mol Cell. 28: 1029–1044.

Dunleavy, E.M., Roche, D., Tagami, H., Lacoste, N., Ray-Gallet, D., Nakamura, Y., et al. (2009) HJURP Is a Cell-Cycle-Dependent Maintenance and Deposition Factor of CENP-A at Centromeres. Cell. 137: 485–497.

Filipescu, D., Szenker, E., and Almouzni, G. (2013) Developmental roles of histone H3 variants and their chaperones. Trends Genet. 29: 630–640.

Finn, R.M., Ellard, K., Eirín-López, J.M., and Ausió, J. (2012) Vertebrate nucleoplasmin and NASP: Egg histone storage proteins with multiple chaperone activities. FASEB J. 26: 4788–4804.

Foltz, D.R., Jansen, L.E.T., Bailey, A.O., Yates, J.R., Bassett, E.A., Wood, S., et al. (2009) Centromere-Specific Assembly of CENP-A Nucleosomes Is Mediated by HJURP. Cell. 137: 472–484.

Le Goff, S., Keçeli, B.N., Jeřábková, H., Heckmann, S., Rutten, T., Cotterell, S., et al. (2020) The H3 histone chaperone NASP SIM3 escorts CenH3 in Arabidopsis. Plant J. 101: 71–86.

Henikoff, S., and Dalal, Y. (2005) Centromeric chromatin: What makes it unique? Curr Opin Genet Dev. 15: 177–184.

Hirsch, C.D., Wu, Y., Yan, H., and Jiang, J. (2009) Lineage-specific adaptive evolution of the centromeric protein cenh3 in diploid and allotetraploid oryza species. Mol Biol Evol. 26: 2877–2885.

Ingouff, M., Rademacher, S., Holec, S., Šoljić, L., Xin, N., Readshaw, A., et al. (2010) Zygotic resetting of the HISTONE 3 variant repertoire participates in epigenetic reprogramming in arabidopsis. Curr Biol. 20: 2137–2143.

Jiang, D., and Berger, F. (2017) Histone variants in plant transcriptional regulation. Biochim Biophys Acta - Gene Regul Mech. 1860: 123–130.

Karimi-Ashtiyani, R., Ishii, T., Niessen, M., Stein, N., Heckmann, S., Gurushidze, M., et al. (2015) Point mutation impairs centromeric CENH3 loading and induces haploid plants. Proc Natl Acad Sci U S A. 112: 11211–11216.

Kawashima, T., Lorković, Z.J., Nishihama, R., Ishizaki, K., Axelsson, E., Yelagandula, R., et al. (2015) Diversification of histone H2A variants during plant evolution. Trends Plant Sci. 20: 419–425.

Kawashima, T., Maruyama, D., Shagirov, M., Li, J., Hamamura, Y., Yelagandula, R., et al. (2014) Dynamic F-actin movement is essential for fertilization in Arabidopsis thaliana. Elife. 3: 1–18.

Khorasanizadeh, S. (2004) The Nucleosome. Cell. 116: 259–272.

Kuppu, S., Ron, M., Marimuthu, M.P.A., Li, G., Huddleson, A., Siddeek, M.H., et al. (2020) A variety of changes, including CRISPR/Cas9-mediated deletions, in CENH3 lead to haploid induction on outcrossing. Plant Biotechnol J. 18: 2068–2080.

Kurihara, D., Mizuta, Y., Nagahara, S., and Higashiyama, T. (2021) ClearSeeAlpha: Advanced Optical Clearing for Whole-Plant Imaging. Plant Cell Physiol. 62: 1302–1310.

Kurihara, D., Mizuta, Y., Sato, Y., and Higashiyama, T. (2015) ClearSee: A rapid optical clearing reagent for whole-plant fluorescence imaging. Dev. 142: 4168–4179.

Lermontova, I., Schubert, V., Fuchs, J., Klatte, S., Macas, J., and Schubert, I. (2006) Loading of Arabidopsis centromeric histone CENH3 occurs mainly during G2 and requires the presence of the histone fold domain. Plant Cell. 18: 2443–2451.

Liu, C.P., Jin, W., Hu, J., Wang, M., Chen, J., Li, G., et al. (2021) Distinct histone H3-H4 binding modes of sNASP reveal the basis for cooperation and competition of histone chaperones. Genes Dev. 35: 1610–1624.

Liu, Y., Chen, L., Wang, N., Wu, B., Bao, H., and Huang, H. (2022) Structural basis for histone H3 recognition by NASP in Arabidopsis. J Integr Plant Biol.

Liu, Z., Chen, O., Wall, J.B.J., Zheng, M., Zhou, Y., Wang, L., et al. (2017) Systematic comparison of 2A peptides for cloning multi-genes in a polycistronic vector. Sci Rep. 7: 1–9.

Luger, K., Mä Der, A.W., Richmond, R.K., Sargent, D.F., and Richmond, T.J. (1997) Luger K, Mäder AW, Richmond RK, Sargent DF, Richmond TJ. Crystal structure of the nucleosome core particle at 2.8 A resolution. Nature. 389: 251–260.

Maheshwari, S., Ishii, T., Brown, C.T., Houben, A., and Comai, L. (2017) Centromere location in Arabidopsis is unaltered by extreme divergence in CENH3 protein sequence. Genome Res. 27: 471–478.

Maheshwari, S., Tan, E.H., West, A., Franklin, F.C.H., Comai, L., and Chan, S.W.L. (2015) Naturally Occurring Differences in CENH3 Affect Chromosome Segregation in Zygotic Mitosis of Hybrids. PLoS Genet. 11: 1–20.

Maksimov, V., Nakamura, M., Wildhaber, T., Nanni, P., Ramström, M., Bergquist, J., et al. (2016) The H3 chaperone function of NASP is conserved in Arabidopsis. Plant J. 88: 425–436.

Malik, H.S., and Henikoff, S. (2009) Major Evolutionary Transitions in Centromere Complexity. Cell. 138: 1067–1082.

Malik, H.S., Vermaak, D., and Henikoff, S. (2002) Recurrent evolution of DNA-binding motifs in the Drosophila centromeric histone. Proc Natl Acad Sci U S A. 99: 1449–1454.

Marimuthu, M.P.A., Maruthachalam, R., Bondada, R., Kuppu, S., Tan, E.H., Britt, A., et al. (2021) Epigenetically mismatched parental centromeres trigger genome elimination in hybrids. Sci Adv. 7: 1–19.

Mayo, K.J., Gonzales, B.J., Mason, H.S. (2006) Genetic transformation of tobacco NT1 cells with *Agrobacterium tumefaciens*. Nat Protoc. 1: 1105–1111.

Medina□Pritchard, B., Lazou, V., Zou, J., Byron, O., Abad, M.A., Rappsilber, J., et al. (2020) Structural basis for centromere maintenance by Drosophila CENP LA chaperone CAL 1. EMBO J. 39: 1–21.

Min, Y., Frost, J.M., and Choi, Y. (2019) Nuclear Chaperone ASF1 is Required for Gametogenesis in Arabidopsis thaliana. Sci Rep. 9: 13959.

Morris, C.A., and Moazed, D. (2007) Centromere Assembly and Propagation. Cell. 128: 647–650.

Nagaki, K., Terada, K., Wakimoto, M., Kashihara, K., and Murata, M. (2010) Centromere targeting of alien CENH3s in arabidopsis and tobacco cells. Chromosom Res. 18: 203–211.

Naish, M., Alonge, M., Wlodzimierz, P., Tock, A.J., Abramson, B.W., Schmücker, A., et al. (2021) The genetic and epigenetic landscape of the Arabidopsis centromeres. Science. 374: eabi7489.

Osakabe, A., Tachiwana, H., Matsunaga, T., Shiga, T., Nozawa, R.S., Obuse, C., et al. (2010) Nucleosome formation activity of human somatic Nuclear Autoantigenic Sperm Protein (sNASP). J Biol Chem. 285: 11913–11921.

Palladino, J., Chavan, A., Sposato, A., Mason, T.D., and Mellone, B.G. (2020) Targeted De Novo Centromere Formation in Drosophila Reveals Plasticity and Maintenance Potential of CENP-A Chromatin. Dev Cell. 52: 379–394.e7.

Phansalkar, R., Lapierre, P., and Mellone, B.G. (2012) Evolutionary insights into the role of the essential centromere protein CAL1 in Drosophila. Chromosom Res. 20: 493–504.

Pusarla, R.H., and Bhargava, P. (2005) Histones in functional diversification: Core histone variants. FEBS J. 272: 5149–5168.

Ravi, M., and Chan, S.W.L. (2010) Haploid plants produced by centromere-mediated genome elimination. Nature. 464: 615–618.

Ravi, M., Kwong, P.N., Menorca, R.M.G., Valencia, J.T., Ramahi, J.S., Stewart, J.L., et al. (2010) The rapidly evolving centromere-specific histone has stringent functional requirements in Arabidopsis thaliana. Genetics. 186: 461–471.

Ravi, M., Marimuthu, M.P.A., Tan, E.H., Maheshwari, S., Henry, I.M., Marin-Rodriguez, B., et al. (2014) A haploid genetics toolbox for Arabidopsis thaliana. Nat Commun. 5: 1–8.

Richardson, R.T., Alekseev, O.M., Grossman, G., Widgren, E.E., Thresher, R., Wagner, E.J., et al. (2006) Nuclear autoantigenic sperm protein (NASP), a linker histone chaperone that is required for cell proliferation. J Biol Chem. 281: 21526–21534.

Richardson, R.T., Batova, I.N., Widgren, E.E., Zheng, L.X., Whitfield, M., Marzluff, W.F., et al. (2000) Characterization of the histone H1-binding protein, NASP, as a cell cycle-regulated somatic protein. J Biol Chem. 275: 30378–30386.

Rosin, L., and Mellone, B.G. (2016) Co-evolving CENP-A and CAL1 Domains Mediate Centromeric CENP-A Deposition across Drosophila Species. Dev Cell. 37: 136–147.

Rosin, L.F., and Mellone, B.G. (2017) Centromeres Drive a Hard Bargain. Trends Genet. 33: 101–117.

Roure, V., Medina-Pritchard, B., Lazou, V., Rago, L., Anselm, E., Venegas, D., et al. (2019) Reconstituting Drosophila Centromere Identity in Human Cells. Cell Rep. 29: 464–479.e5.

Sanei, M., Pickering, R., Kumke, K., Nasuda, S., and Houben, A. (2011) Loss of centromeric histone H3 (CENH3) from centromeres precedes uniparental chromosome elimination in interspecific barley hybrids. Proc Natl Acad Sci U S A. 108: E498–505.

She, W., Grimanelli, D., Rutowicz, K., Whitehead, M.W.J., Puzio, M., Kotliński, M., et al. (2013) Chromatin reprogramming during the somatic-to-reproductive cell fate transition in plants. Dev. 140: 4008–4019.

Stoler, S., Rogers, K., Weitze, S., Morey, L., Fitzgerald-Hayes, M., and Baker, R.E. (2007) Scm3, an essential Saccharomyces cerevisiae centromere protein required for G2/M progression and Cse4 localization. Proc Natl Acad Sci U S A. 104: 10571–10576.

Takeuchi, H., and Higashiyama, T. (2016) Tip-localized receptors control pollen tube growth and LURE sensing in Arabidopsis. Nature. 531: 245–248.

Talbert, P.B., and Henikoff, S. (2010) Histone variants ancient wrap artists of the epigenome. Nat Rev Mol Cell Biol. 11: 264–275.

Talbert, P.B., and Henikoff, S. (2017) Histone variants on the move: Substrates for chromatin dynamics. Nat Rev Mol Cell Biol. 18: 115–126.

Talbert, P.B., Masuelli, R., Tyagi, A.P., Comai, L., and Henikoff, S. (2002) Centromeric localization and adaptive evolution of an Arabidopsis histone H3 variant. Plant Cell. 14: 1053–1066.

Tsutsui, H., and Higashiyama, T. (2017) pKAMA-ITACHI vectors for highly efficient CRISPR/Cas9-mediated gene knockout in Arabidopsis thaliana. Plant Cell Physiol. 58: 46–56.

Yuan, J., Guo, X., Hu, J., Lv, Z., and Han, F. (2015) Characterization of two CENH3 genes and their roles in wheat evolution. New Phytol. 206: 839–851.

Zhang, W., Lee, H.R., Koo, D.H., and Jiang, J. (2008) Epigenetic modification of centromeric chromatin: Hypomethylation of DNA sequences in the CENH3-associated chromatin in Arabidopsis thaliana and maize. Plant Cell. 20: 25–34.

Zhong, Z., Wang, Y., Wang, M., Yang, F., Thomas, Q.A., Xue, Y., et al. (2022) Histone chaperone ASF1 mediates H3.3-H4 deposition in Arabidopsis. Nat Commun. 13.

Zhou, Z., Feng, H., Zhou, B.R., Ghirlando, R., Hu, K., Zwolak, A., et al. (2011) Structural basis for recognition of centromere histone variant CenH3 by the chaperone Scm3. Nature. 472: 234–238.

Zhu, Y., Weng, M., Yang, Y., Zhang, C., Li, Z., Shen, W.H., et al. (2011) Arabidopsis homologues of the histone chaperone ASF1 are crucial for chromatin replication and cell proliferation in plant development. Plant J. 66: 443–455.

