## Supplementary Fig. S for "The Chaperone NASP Contributes to *De Novo* Deposition of the Centromeric Histone Variant CENH3 in *Arabidopsis* Early Embryogenesis"

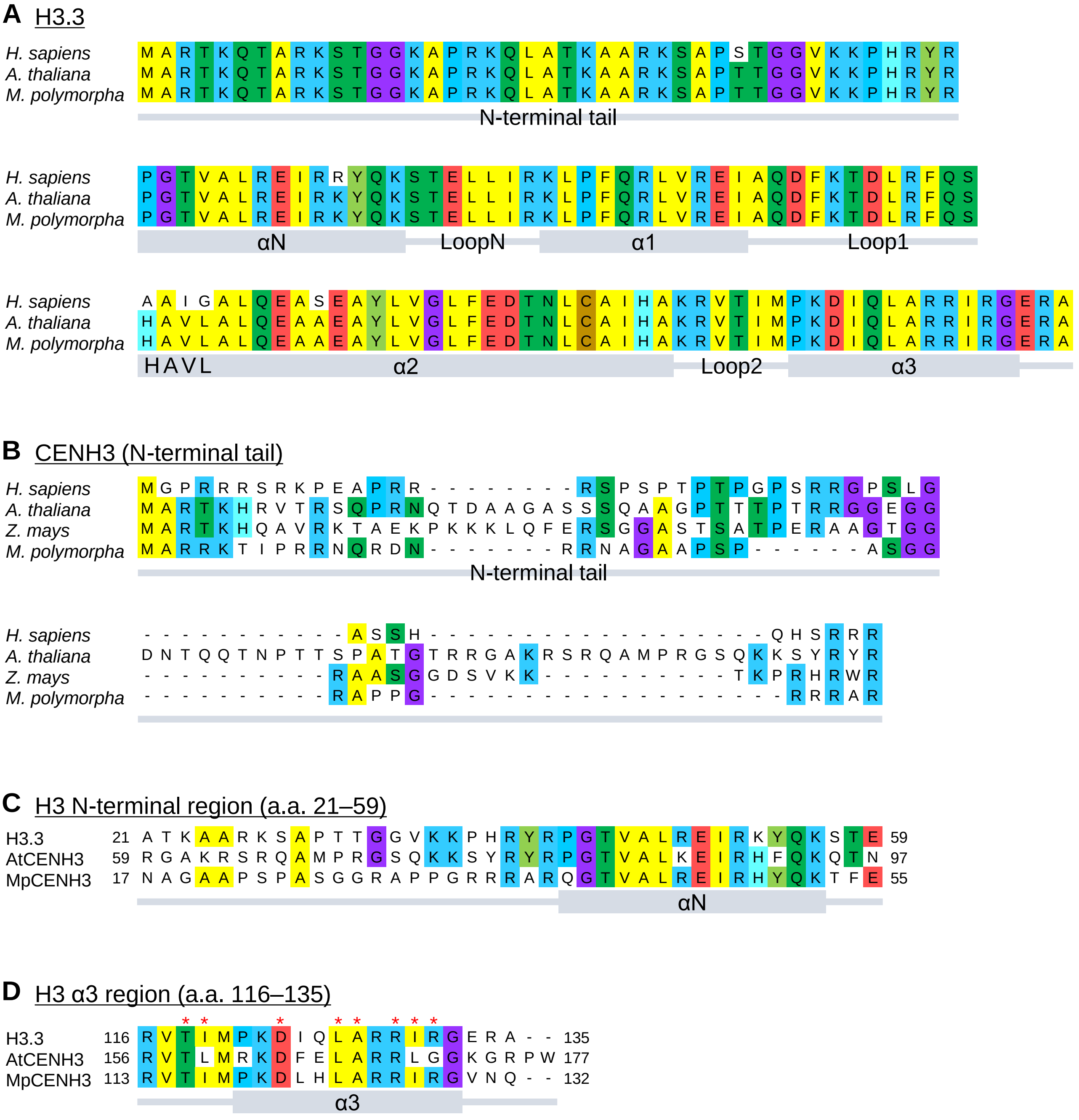


**Supplementary Fig. S1**

Sequence alignment of histone H3.3 and N-terminal tail of CENH3, related to Fig. 1A. Protein secondary structure is indicated below the alignment. (A) Full-length histone H3.3 from *H. sapiens*, *A. thaliana*, and *M. polymorpha*. (B) N-terminal tail of CENH3 from *H. sapiens*, *A. thaliana*, *Z. mays*, and *M. polymorpha*. (C, D) N-terminal region (a.a. 21–59) and α3 region (a.a. 116–135), which correspond to two sets of the AtNASP-binding region to canonical H3 (Liu et al. 2022), of *A. thaliana*/*M. polymorpha* H3.3, *A. thaliana* CENH3, and *M. polymorpha* CENH3. Red asterisks in (D) indicate corresponding residues involved in the interaction between H3 and AtNASP according to the crystal structure (Liu et al. 2022).

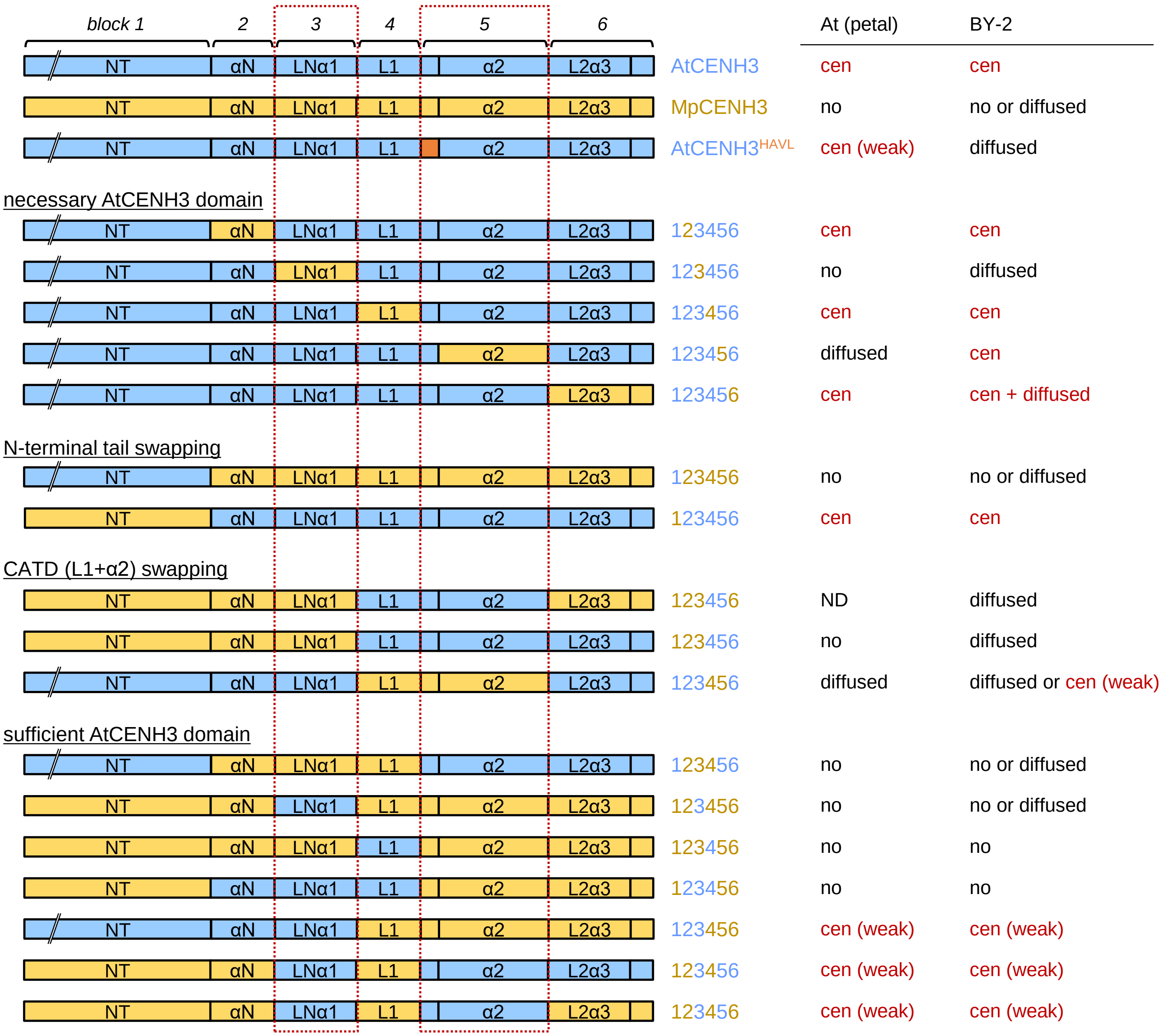


**Supplementary Fig. S2**

A summary of the localization assay, related to Fig. 1 and Supplementary Fig. S3. CENH3 regions of *A. thaliana* and *M. polymorpha* are shown with light blue and yellow, respectively. Assessments for the localization of each chimeric CENH3 variant are shown based on observation as follows: cen, dot-like centromere-focused signal; cen (weak), dot-like centromere-focused signal despite weak intensity; diffused, diffused signal in the entire nucleus; no, undetectable signal in the nucleus; ND, no data.

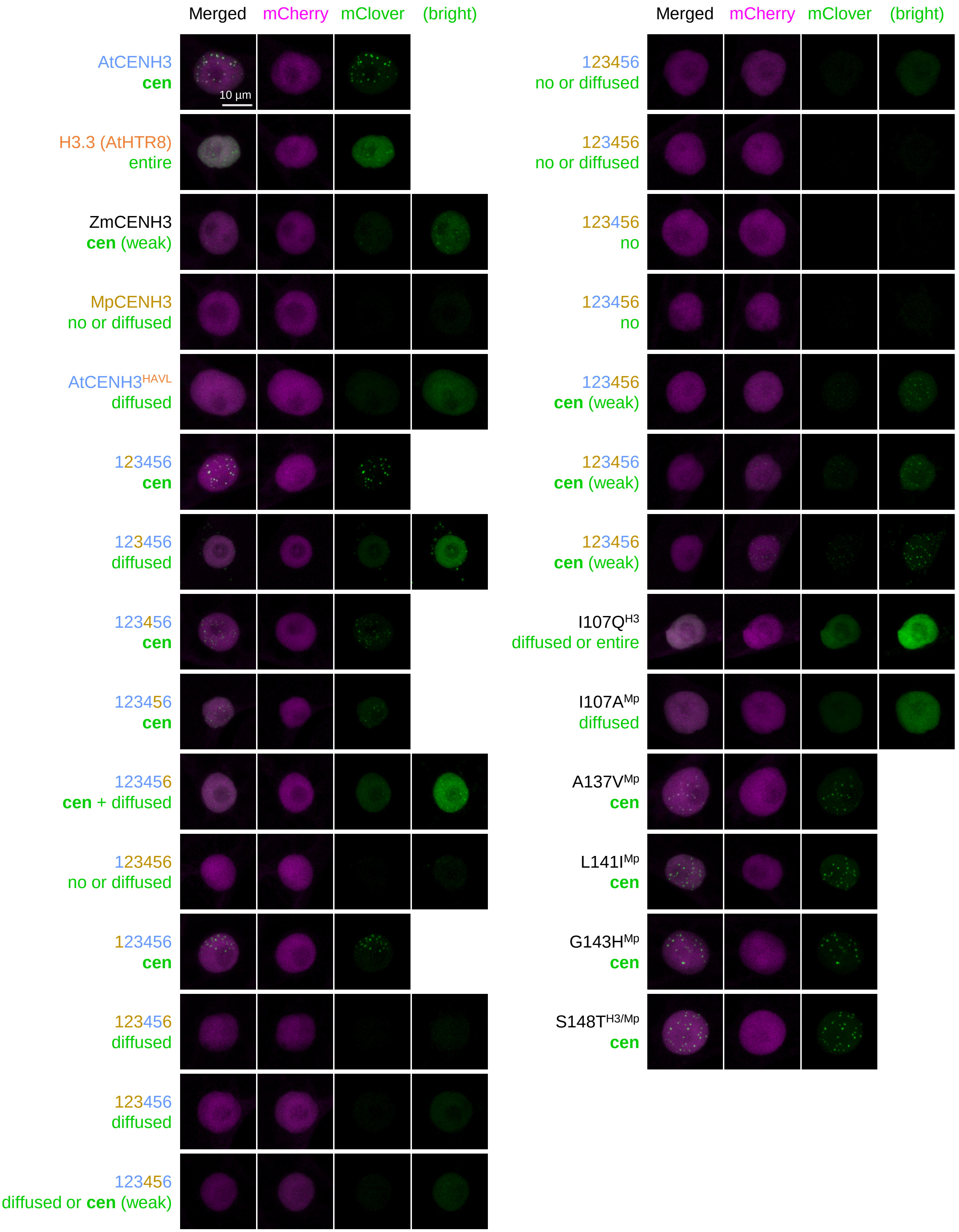


**Supplementary Fig. S3**
Representative images of tobacco BY-2 cultured cell lines expressing each chimeric CENH3 variant. Note that mClover-fused CENH3 variant showed different localization among lines, while signals of co-translated NLS-mCherry are comparable. For cell lines with weaker mClover signal, images were processed to be high-contrast images (as “bright”).
